# HOPS recognizes each SNARE, assembling ternary *trans*-complexes for sudden fusion upon engagement with the 4th SNARE

**DOI:** 10.1101/751404

**Authors:** Hongki Song, Amy Orr, Max Harner, William Wickner

## Abstract

Vacuole fusion requires SNAREs, Sec17/18, a Rab, and HOPS. We find that co-incubation of HOPS, proteoliposomes bearing the Rab and R-SNARE, and proteoliposomes with the Rab and any two Q-SNAREs yields a *trans* complex which includes these 3 SNAREs. The missing Q-SNARE then triggers a burst of fusion, indicating that each HOPS, R-, and QxQy-SNARE *trans*-complex is an activated intermediate for functional Qz-SNARE incorporation. HOPS can assemble activated fusion intermediates because it recognizes each of the four SNAREs, binding them independently. HOPS-dependent fusion is saturable for each Q-SNARE, indicating saturable functional sites on HOPS. Though a nonspecific tether allows fusion with pre-assembled Q-SNAREs, only HOPS catalyzes fusion when the Q-SNAREs are not pre-assembled by ushering each Q-SNARE into a functional complex. In contrast, there is little spontaneous functional assembly of the 3 Q-SNAREs. HOPS thus recognizes each of the 4 SNAREs to assemble a versatile set of activated fusion intermediates.

## Introduction

Membrane fusion at each organelle is orchestrated by conserved families of proteins and lipids with complex binding relationships (Wickner and Rizo, 2017). Tethering effectors bind Rab family GTPases to hold membranes in apposition (Baker and Hughson, 2016). SNARE (soluble N-ethylmaleimide-sensitive-factor attachment receptor) proteins are found on both fusion partners, either in *cis*-SNARE complexes if all are anchored to one membrane or *trans*-SNARE complexes if anchored to two apposed membranes. SNAREs have heptad-repeat SNARE domains of approximately 50 aminoacyl residues with a central arginyl (R) or glutaminyl (Q) residue. SNAREs are grouped by sequence homology into four families, R, Qa, Qb, and Qc (Fasshauer et al., 1998). SNARE complexes have one member each of the R, Qa, Qb, and Qc families, with their α-helical SNARE domains wrapped together in a coiled coil (Sutton et al., 1998). This 4-SNARE bundle is stabilized by the interior disposition of apolar residues, with the exception of 1 arginyl and 3 glutaminyl residues in the center of the SNARE domain, termed the 0-layer. SNARE complex assembly can be promoted by Sec1/Munc18 (SM) family proteins (Baker et al., 2015; Jiao et al., 2018; Rizo and Südhof, 2012). Disassembly of the post-fusion *cis*-SNARE complexes is catalyzed by the ATP-driven chaperone Sec18/NSF, stimulated by its co-chaperone Sec17/αSNAP (White et al., 2018). Sec17 and Sec18 also function earlier, stimulating the fusion of docked membranes (Song et al., 2017; Zick et al., 2015). Fusion also requires fatty acyl fluidity (Zick and Wickner, 2016), acidic lipids and phosphoinositides to promote the binding of peripheral membrane fusion proteins (Cheever et al., 2001; Mima and Wickner, 2009; Orr et al., 2015), and nonbilayer-prone lipids (Zick et al., 2014) to enable the bilayer rearrangements which are essential for fusion.

We study membrane fusion mechanisms with the vacuole (lysosome) of *S. cerevisiae*. Vacuoles undergo constant fission and fusion, regulated by growth medium osmolarity. Mutations which block fusion allow continued fission, resulting in a visibly altered **<u>va</u>**cuole **<u>m</u>**orphology which allowed selection of *vam* mutants in fusion (Wada et al., 1992). The *VAM* genes encode proteins which are unique to vacuole fusion: the Rab Ypt7, the 6 subunits of the HOPS (**<u>ho</u>**motypic fusion and vacuole **<u>p</u>**rotein **<u>s</u>**orting) tethering and SM complex (Nakamura et al., 1997; Seals et al., 2000; Wurmser et al., 2000), and the R, Qa, and Qc SNAREs of this organelle (hereafter referred to as R, Qa, and Qc). Other vacuole fusion proteins such as the Qb SNARE, Sec17, and Sec18 are also required in the exocytic pathway and were not identified in the *vam* screen since their loss is lethal.

Vacuole fusion has been extensively studied *in vivo*, *in vitro* with the purified organelle, and as reconstituted with proteoliposomes bearing all-purified components (Mima et al., 2008; Zick and Wickner, 2016). The “priming” stage of fusion, which precedes organelle association, entails phosphoinositide synthesis (Mayer et al., 2000) and Sec17- and Sec18-dependent *cis*-SNARE complex disassembly (Mayer et al., 1996). Priming is a prerequisite for tethering (Mayer, 1997), which is largely mediated by the affinities of two of the HOPS subunits (Vps39 and Vps41) for the Rab Ypt7 on each vacuole membrane (Brett et al., 2008). Vacuoles also have a “back-up” tethering system through the affinity of the PX domain of the Qc SNARE for PtdIns3P in *trans* (Zick and Wickner, 2014). HOPS has been proposed to catalyze the productive association in *trans* of R-SNAREs with the Qa-SNARE from the closely apposed membrane, initiating the formation of a *trans* 4-SNARE complex (Baker et al., 2015). Fusion can be supported by HOPS and SNAREs alone, but is further accelerated by Sec17 and Sec18p without requiring ATP hydrolysis (Song et al., 2017).

These fusion proteins and lipids show interdependent co-enrichment on docked vacuoles at a ring-shaped microdomain surrounding the directly apposed bilayers (Wang et al., 2002; Fratti et al., 2004). The full panoply of affinities and functional interactions of these fusion components is only now emerging. SM proteins are known to bind to Qa SNAREs, and a conserved R-SNARE binding site has been found on the Vps33 subunit of HOPS and on other SM proteins (Baker et al., 2015). Other subunits of HOPS, Vps16 and Vps18, bind the Qc SNARE (Krämer and Ungermann, 2011) through the distinct PX region of Qc that is N-terminal to the Qc SNARE domain (Stroupe et al., 2006). Direct affinity of HOPS for Qb has not been reported. However, in a chemically-defined subreaction of fusion, proteoliposomes bearing Ypt7 and the R-SNARE underwent HOPS-dependent assembly of all of the Q-SNAREs, including Qb, into a 4-SNARE complex (Orr et al., 2017). Once a 4-SNARE complex has assembled, several Sec17/αSNAP molecules can bind along its length (Zhao et al., 2015). The N-terminal apolar loop of SNARE-bound Sec17 has direct affinity for the lipid bilayer (Zick et al., 2015), while the membrane-distal C-terminus binds Sec18/NSF (Marz et al., 2003; Winter et al., 2009). HOPS also has direct affinity for phosphoinositides such as PtdIns3P (Stroupe et al., 2006) and for acidic lipids (Karunakaran and Wickner, 2013).

The availability of pure and active fusion proteins and their reconstitution into model subreactions has allowed the detection of additional functional affinities among these components. We now report that HOPS supports the assembly of an active state between membranes bearing Ypt7 and the R-SNARE and other membranes bearing Ypt7 and any two of the 3 Q-SNAREs. HOPS and these 3 SNAREs are in a stable, isolable *trans* complex. Upon encountering the third Q-SNARE, this active state supports strikingly rapid fusion. Without HOPS, proteoliposomes with any two Q-SNAREs are extremely slow to assemble with the third. The capacity of HOPS to form 3-SNARE rapid fusion intermediates is supported by the finding that HOPS binds directly to each SNARE. As a complementary demonstration of the functionality of HOPS recognition of each Q-SNARE, we show that the HOPS-mediated fusion of R- and single Q-SNARE proteoliposomes is saturable over a wide range of concentations of soluble forms of the other Q-SNAREs, whereas there is no saturation when HOPS is replaced by an artificial tether. Thus HOPS recognition of each Q-SNARE supports its functional assembly with the others without an obligate order.

## Results

In detergent micellar solution, SNAREs spontaneously assemble into 4-SNARE complexes or subcomplexes (Fukuda et al., 2000). In the context of lipid bilayers, *trans*-SNARE complex assembly may be affected by the membrane anchoring of SNAREs, by membrane apposition through tethering, and by the affinity of HOPS for the SNAREs on each membrane. Tethering per se will support functional trans-SNARE formation between R- and 3Q-SNARE proteoliposomes (Song and Wickner, 2019); does it suffice if the Q-SNAREs are not pre-assembled?

### HOPS-mediated assembly of fusion intermediates

To study the functional intermediates in SNARE complex assembly, we assayed fusion without an added tether, with tethering by the physiological and multifunctional HOPS complex bound to the Rab Ypt7 on each membrane, or with a simple synthetic tether. Our synthetic tether consists of dimeric glutathione S-transferase (GST) fused to a PX domain that can bind to PtdIns3P in each proteoliposomal membrane (Song and Wickner, 2019).

Proteoliposomes bearing Ypt7 and R-SNARE with lumenally entrapped biotinylated phycoerythrin were mixed with proteoliposomes bearing Ypt7 and the 3 Q-SNAREs with entrapped Cy5-streptavidin. These mixed proteoliposomes were incubated without tethering agent, with HOPS, or with GST-PX. Either tethering agent sufficed for fusion, which was detected by the FRET from the mixing of the lumenal dyes (Figure 1A). Similar proteoliposomes in which the Qc SNARE bore a synthetic C-terminal anchor (Xu and Wickner, 2012) showed similar fusion (Figure 1B). Though tethering is required, this shows that productive association of the three pre-assembled Q-SNAREs and the R-SNARE into functional *trans*-SNARE complex does not always require catalysis by the Vps33 SM-protein subunit of HOPS. This is consistent with earlier findings that deletion within the R and Qa SNARE recognition domains of the Vps33 subunit of HOPS still permits fusion between R- and QaQbQc-SNARE proteoliposomes (Baker et al., 2015). To explore the role of HOPS in the assembly of each Q-SNARE into functional *trans*-SNARE subcomplexes, Ypt7/2Q-SNARE proteoliposomes were prepared in 3 batches, termed QbQc_tm_, QaQc_tm_, and QaQb, each bearing Ypt7 and 2 membrane-anchored Q-SNAREs and omitting Qa, Qb, or Qc, respectively. Each of these Ypt7/2Q-SNARE proteoliposomes fused well with Ypt7/R-SNARE proteoliposomes when incubated with both HOPS and the soluble form (without membrane anchor) of the omitted Q-SNARE (Figure 1C-E, blue curve). Both HOPS and the soluble SNARE were essential for fusion in each case (no HOPS, black curve; no soluble SNARE, green curve). HOPS provided more than just a tethering function, since the GST-PX synthetic tether gave almost no fusion (GST-PX and soluble SNARE, red curve), though it had sufficed when all three of the Q-SNAREs were pre-assembled (A and B, red curve). Thus HOPS allows rapid and efficient recruitment of any Q-SNARE and does not merely do so through its tethering function. Without HOPS, there is little 3Q complex assembly on 2Q-SNARE proteoliposomes which had been incubated with the third Q-SNARE. Had such assembly occurred, the resulting 3Q-SNARE proteoliposomes would have been capable of GST-PX mediated fusion (Figure 1A, B). With HOPS present, docking, SNARE complex assembly, and fusion begins without lag (Figure 1C-E).

**Figure 1.**
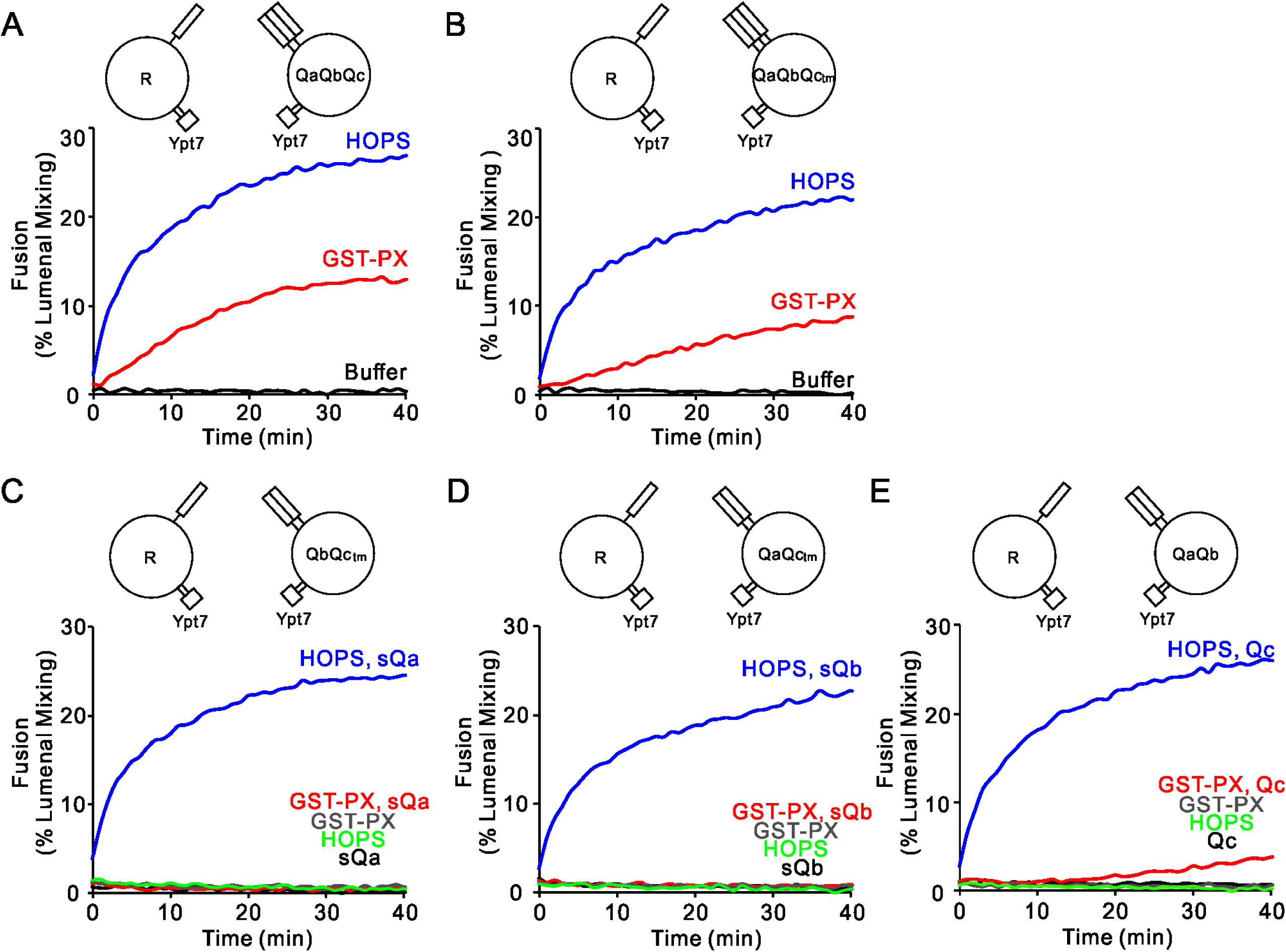
HOPS recruits each Q-SNARE, whereas a simple tether (GST-PX) does not. **(A-E)** Fusion reactions had proteoliposomes bearing either R- or Q-SNARE combinations as indicated at 1:16000 SNARE:lipid molar ratio. Fusion was assayed between R and **(A)** QaQbQc, **(B)** QaQbQc_tm_, **(C)** QbQc_tm_, **(D)** QaQc_tm_, or **(E)** QaQb proteoliposomes. Fusion reactions had 500nM GST-PX, 50nM HOPS and 4μM soluble SNAREs as indicated. All proteoliposomes had Ypt7-tm at a 1:8,000 protein: lipid molar ratio. Kinetic curves of content mixing assays in this figure are representative of n ≥ 3 experiments; average and standard deviations of fusion from three independent experiments are in Figure 1-figure supplement 1.

### Limited spontaneous Q-SNARE assembly

Since wild-type Qa and Qb are membrane anchored, and only Qc is soluble, we analyzed the fusion of R- and QaQb-proteoliposomes with soluble Qc in more depth. The full kinetic time course shows that the synthetic tether GST-PX did support detectable fusion of R- and QaQb-proteoliposomes with added Qc, but only very slowly and after a 10-minute lag (Figure 1E and Figure 1-figure supplement 2, curve 1). This lag was eliminated and the fusion rate enhanced by a 30 minute preincubation of the QaQb-SNARE proteoliposomes with Qc-SNARE prior to addition of the GST-PX tether (Figure 1-figure supplement 2, curves 1 vs. 3 and 7), revealing a capacity for slow spontaneous assembly of stable 3Q-SNARE complex from QaQb-proteoliposomes and Qc (curve 7). This was not seen with GST-PX and 2Q-SNARE proteoliposomes lacking Qa or Qb and supplemented with sQa or sQb, respectively (Figure 1C, D), suggesting that these assembly events are either kinetically too slow or thermodynamically unfavorable.

As a second, complementary assay for spontaneous assembly of functional 3Q-SNARE complex, we employed the R-SNARE without its membrane anchor, termed soluble-R (sR), a known fusion inhibitor (Thorngren et al., 2004; Zick and Wickner, 2014; Song and Wickner, 2019). Inhibition by sR can employ two mechanisms: 1. sR may compete for the conserved R-SNARE binding groove on the Vps33 subunit of HOPS (Baker et al., 2015). 2. If a stable SNARE complex is assembled which includes the sR-SNARE, subsequent fusion with R-SNARE proteoliposomes is blocked; for example, preincubation of Ypt7/3Q- and Ypt7/R-proteoliposomes with sR for 30 min prior to HOPS addition blocks their fusion (Zick and Wickner, 2014). R- and QbQc_tm_ or QaQc_tm_-SNARE proteoliposomes were mixed and preincubated with or without sR and with or without the third sQ for 30 minutes, then HOPS and (where absent) the sQ were added to initiate fusion (Figure 1-figure supplement 3, A and B). The fusion seen without sR (curves e, f) was inhibited approximately 2-fold by sR (curves a-d) without regard to the order of addition and incubation, which may reflect sR competition for a conserved site on Vps33. However, there was complete fusion inhibition when QaQb-proteoliposomes were preincubated with both sR and Qc for 30 min prior to HOPS addition (C, curve a), suggesting sRQaQbQc assembly. The contrast between the full inhibition by sR when preincubated with Qc and QaQb proteoliposomes (Figure 1-figure supplement 3, C, curve a) and the lack of inhibition enhancement when either soluble Qa or soluble Qb is preincubated with sR and QbQc_tm_ or QaQc_tm_ proteoliposomes, respectively (A and B, curve a), is another indication that soluble Qa and Qb do not spontaneously enter into complex with the other Q-SNAREs prior to HOPS addition. In sum, HOPS is required for any one of the Q-SNAREs to assemble rapidly with the others into functional SNARE complex for fusion, though a very slow spontaneous assembly of Qc can occur.

### HOPS-activated R- and 2Q-SNARE assembly intermediates

Since Qc is the one physiologically soluble Q-SNARE, we sought to physically and functionally measure any HOPS stabilized fusion-competent assemblies in *trans*. Ypt7/R-SNARE proteoliposomes were incubated with Ypt7/QaQb proteoliposomes in the presence or absence of HOPS or Qc. Fusion required both HOPS and Qc (Figure 2A, curve d; other incubations lacked HOPS or Qc or both). After 30min of incubation of mixed proteoliposomes with HOPS, the addition of Qc (curve e) triggered abrupt fusion which was not seen without Qc (curve c), showing that a highly active fusion intermediate had accumulated. Samples from each incubation were withdrawn at 33 minutes, solubilized in detergent with an excess of GST-R to competitively block any wild-type R which might have otherwise associated with Qa in the extract, and assayed by pulldown with antibody to Qa to determine the untagged R-SNARE which had become associated with the Qa-SNARE (Figure 2, B and C). While there was background association of the R and Qa SNAREs in incubations without HOPS or Qc (lane a) and maximal association with both HOPS and Qc (lane d), HOPS promoted substantial *trans* complex assembly between R and Qa SNAREs in the absence of Qc (lane c) while fusion remained blocked (Figure 2A, curve c). The addition of Qc at 30 minutes triggered a sudden burst of fusion (Figure 2A, red curve e) with only a modest increase in *trans*-SNARE complex (Figure 2C, e vs c). HOPS thus assembles the R- and QaQb-SNAREs in *trans*, whether directly with each other in 3-SNARE bundles or with the R- and two Q-SNAREs each associated with common HOPS molecules.

**Figure 2.**
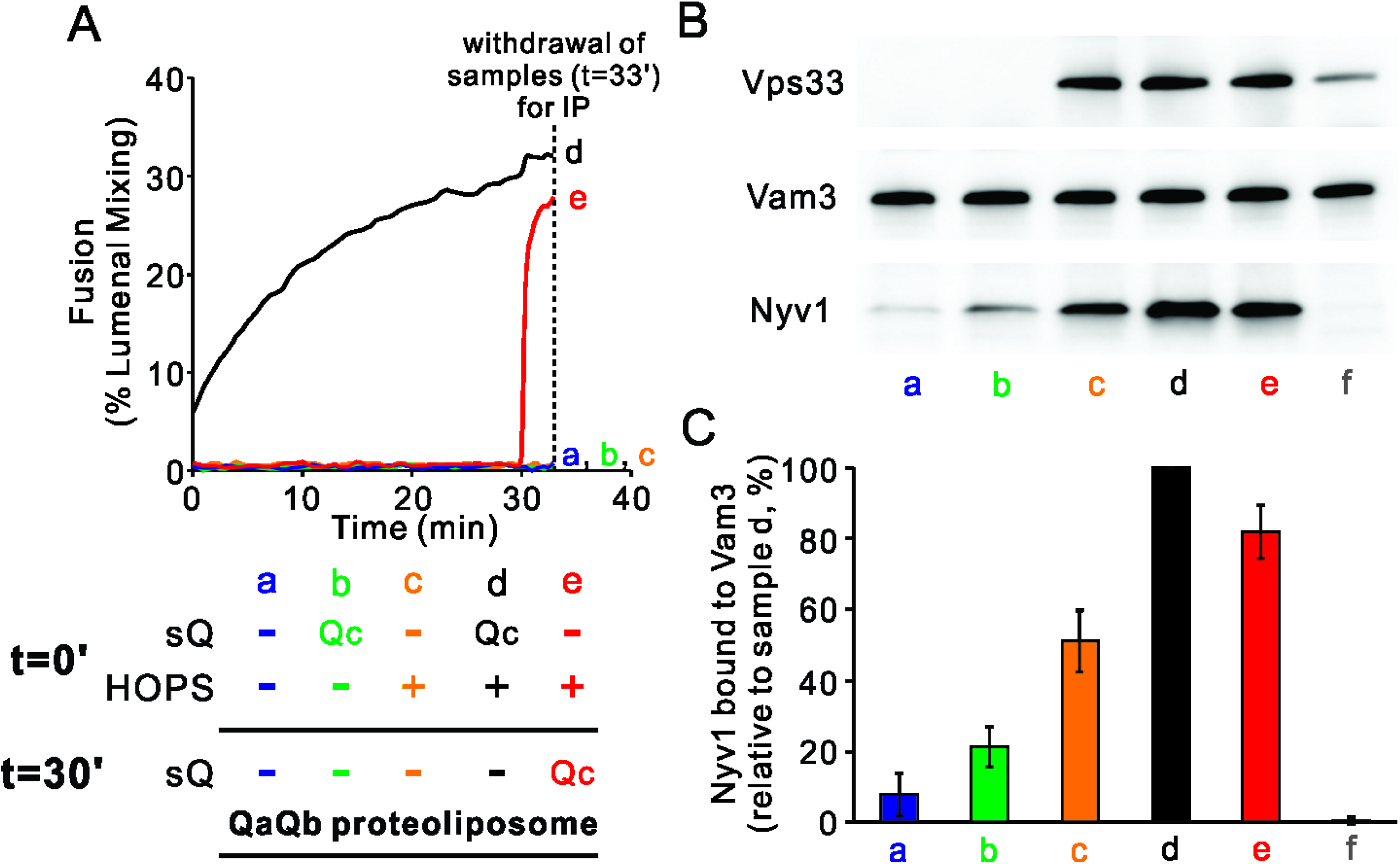
HOPS induces formation of a rapid-fusion intermediate which includes the R- and Qa-SNAREs in *trans* association. Proteoliposomes with Ypt7 and R (1:8000 and 1:16,000 molar ratio to lipids, respectively) were mixed with proteoliposomes with Ypt7 and Qa and Qb SNAREs with 50 nM HOPS and 4 μM Qc where indicated, added either at the start of incubation or after 30min. (A) Fusion was assayed as luminal FRET. After 30min, Qc was added to one sample (e, blue). (B) To measure complex formation, the amount of R SNARE that was immunoprecipitated from a detergent extract with anti-Qa antibody was determined. After incubation for 33 min, samples were placed on ice and mixed with 5 volumes of ice-chilled modified RIPA buffer [20 mM Hepes·NaOH, pH 7.4, 150 mM NaCl, 0.2% (wt/vol) BSA, 1% (vol/vol) Triton X-100,1% (wt/vol) sodium cholate, 0.1% (wt/vol) SDS] containing RIPA buffer washed protein A magnetic beads (ThermoFisher), 5 μM GST-R and 5μg anti-Qa antibody. After the mix was nutated at 4°C for 2 h, beads were washed three times with 1 mL of RIPA buffer. Proteins were eluted with 100μL of SDS sample buffer at 95 °C for 5 min. Eluates were assayed by immunoblot with antibodies to R, Qa and Vps33. (C) Immunoblots for the R-SNARE were scanned from 6 experiments, the band intensity of sample d (HOPS and Qc added at t = 0 min) was set to 100%, and the means and standard deviations are shown.

To determine whether the HOPS-dependent formation of rapid-fusion complex was specific to the Q-SNARE which was last to engage and thereby trigger rapid fusion, Ypt7/R-SNARE proteoliposomes were mixed with each of the three Ypt7/2Q-SNARE proteoliposomes, the soluble Q-SNARE that was not proteoliposome-bound, and HOPS (Figure 3A-C, black curve a). The rate of fusion was compared to that seen when HOPS had been preincubated with the two mixed sets of proteoliposomes without the soluble Q-SNARE for 30min prior to soluble-Q addition (red curve b). For each Q-SNARE, there was distinctly more rapid fusion when the soluble Q-SNARE was added after 30 minutes (Figure 3A-C, red curve b; see Figure 3-figure supplement 1) than when it had been added from the start (curve a), indicating that HOPS allowed the accumulation of fusion-competent intermediates which included either R and QaQb, R and QaQc_tm_, or R and QbQc_tm_, respectively. The formation of the rapid-fusion state required the presence of HOPS during the preincubation (compare curve b to curves c, d). The action of HOPS is not merely due to its tethering function, as the synthetic tether GST-PX does not support fusion at all unless the three Q-SNAREs have been pre-assembled (Figure 1). These assays do not distinguish whether the R- and two Q-SNAREs had entered a 3-helical coiled coils SNARE subcomplex or whether HOPS catalysis consisted of binding the R and two Q-SNAREs in a configuration which could rapidly and functionally receive the third Q-SNARE for fusion.

**Figure 3.**
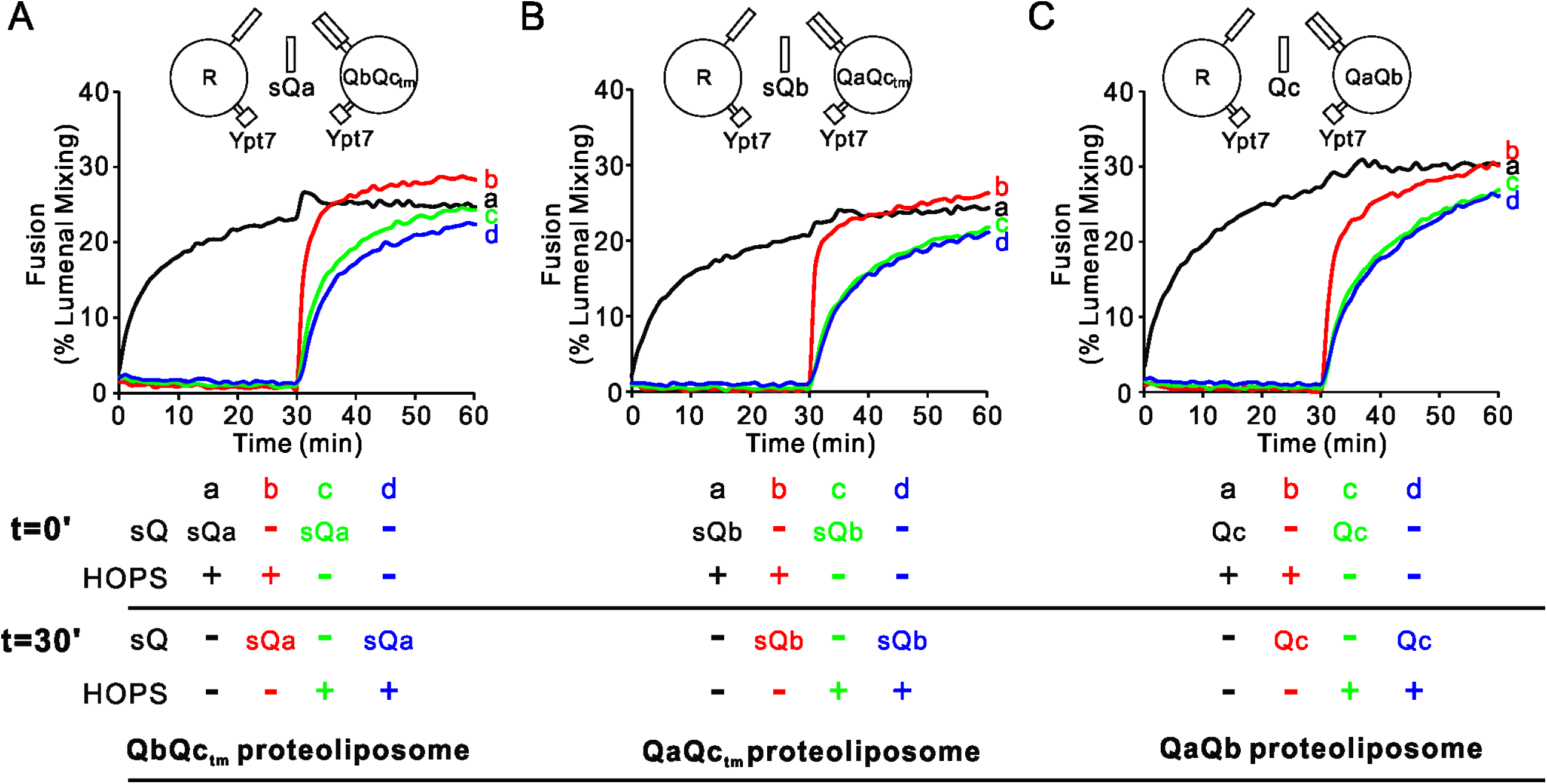
HOPS activates mixed proteoliposomes with R+Ypt7 and 2Q+Ypt7 for a burst of fusion when the third Q-SNARE is supplied. (A-C) Fusion incubations received 50 nM HOPS at t=0 min (a, b) or t=30 min (c, d) with 4μM soluble Q-SNARE at t=0 min (a, c) or t=30 min (b, d) as indicated. Fusion reactions had proteoliposomes bearing either the R mixed with proteoliposomes bearing (A) QbQc_tm_, (B) QaQc_tm_, or (C) QaQb SNAREs at 1:16000 SNARE:lipid molar ratios. All proteoliposomes had Ypt7-tm at a 1:8,000 protein: lipid molar ratio. Content mixing assays in this figure are representative of n ≥ 3 experiments; means and standard deviations for each experiment are presented in Figure 3-figure supplement 1.

### Physical and functional HOPS affinity for each Q-SNARE

These studies show that HOPS can catalyze assembly of a complex which can receive each Q-SNARE for rapid fusion, but do not address whether it has the capacity to bind each SNARE directly. We therefore asked whether HOPS can bind each vacuolar SNARE. Prior studies have shown that HOPS has direct affinity for the PX domain of the Qc SNARE (Stroupe et al., 2006) through its Vps16 and Vps18 subunits (Krämer and Ungermann, 2011) and for the R- and Qa-SNARE domains through its Vps33 SM-family subunit (Baker et al., 2015). HOPS has not been reported to have direct affinity for the Qb SNARE. To evaluate the ability of HOPS to bind to each SNARE, we prepared 6 sets of liposomes, either protein-free liposomes or proteoliposomes bearing one of the four vacuolar SNAREs (including a characterized membrane-anchored form of Qc; Xu and Wickner, 2012) or all four SNAREs. Each set of proteoliposomes was incubated with HOPS, mixed with density medium, overlaid with a density gradient, and subjected to ultracentrifugation. The floated proteolipsomes were assayed by immunoblot for bound HOPS. Though HOPS was not recovered with protein-free liposomes (Figure 4, lane 1; also Figure 4-figure supplement 1), HOPS bound to each of the 4 SNAREs (lanes 2-5) or their complex (lane 6).

**Figure 4.**
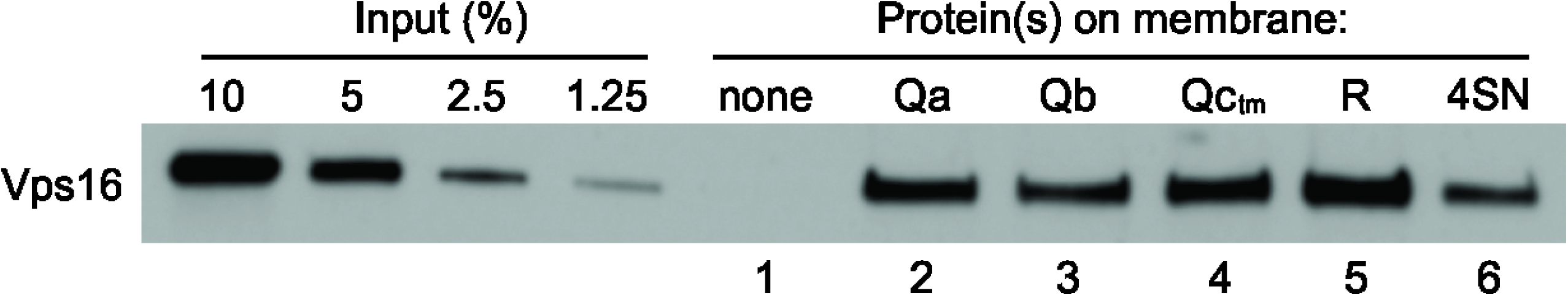
HOPS binds directly to each vacuolar SNARE. PC liposomes with no integral SNAREs, with each individual integrally-bound SNARE, or with all four wild-type SNAREs were incubated with HOPS at a 2-fold molar excess to SNAREs and floated as described. Bound HOPS was analyzed by immunoblot of its Vps16 subunit with a standard curve of the input. Quantification and statistical analysis of HOPS binding from 3 independent experiments is in Figure 4-figure supplement 1.

Since HOPS can bind to each SNARE individually, might it promote the fusion of proteoliposomes bearing a single SNARE, as it does for those with pairs of Q-SNAREs (Figure 1)? We complemented our physical binding assay of HOPS binding to single SNAREs with functional assays of the fusion between proteoliposomes bearing Ypt7 plus the R-SNARE and those with Ypt7 plus a single integrally-anchored Q-SNARE (Figure 5). The fusion of proteoliposomes that have Ypt7, R-SNARE, and lumenally entrapped biotinylated phycoerythrin with those bearing Ypt7, Qa-SNARE, and lumenally entrapped Cy5-streptavidin is supported by sQb, Qc, and an additional agent, either HOPS (Song and Wickner, 2017) or polyethylene glycol (PEG). While HOPS can specifically bind SNAREs, PEG is a nonspecific dehydrating agent (Lentz, 2007) which clusters membranes and promotes SNARE assembly without any SNARE-binding specificity. With PEG, the fusion rate steadily declines with diminishing sQb (Figure 5A, filled bars), as expected for four SNAREs spontaneously assembling into a required tetramer. However, with HOPS the rate is almost constant over the same wide sQb concentration range (Figure 5A, open bars), indicating saturation of an active HOPS binding site for sQb. [Earlier studies of HOPS-mediated fusion between R- and Qa-SNARE proteoliposomes had employed MBP-sQb, and found that it hadn’t exhibited saturable kinetics (Zick and Wickner, 2013). We reproduce this finding (Figure 5-figure supplement 1, filled bars), and note that the MBP “tag” had prevented a high-affinity, saturable engagement with HOPS, which is seen upon proteolytic removal of the tag (open bars).] When Ypt7/R and Ypt7/Qa proteoliposomes were mixed with ample sQb and the concentration of Qc was varied, fusion with PEG tethering was again proportional to the Qc concentration, while fusion with HOPS as the tether showed little change over a wide range of Qc (Figure 5B). This saturation indicates that HOPS has a functional Qc binding site. In a similar approach, proteoliposomes bearing Ypt7 and R were incubated with those bearing Ypt7 and Qb, in the presence of sQa, Qc, and either HOPS or PEG. With HOPS, the rate of fusion was saturable with respect to the concentration of Qa or Qc (Figure 5C, D). HOPS-dependent fusion of Ypt7/R and Ypt7/Qc proteoliposomes, where Qc was fused to a membrane anchor, is also invariant over a wide range of sQa or sQb concentrations (Figure 5E, F); direct comparison with PEG-mediated fusion was not possible, as PEG did not support the SNARE-dependent fusion of these proteoliposomes.

**Figure 5.**
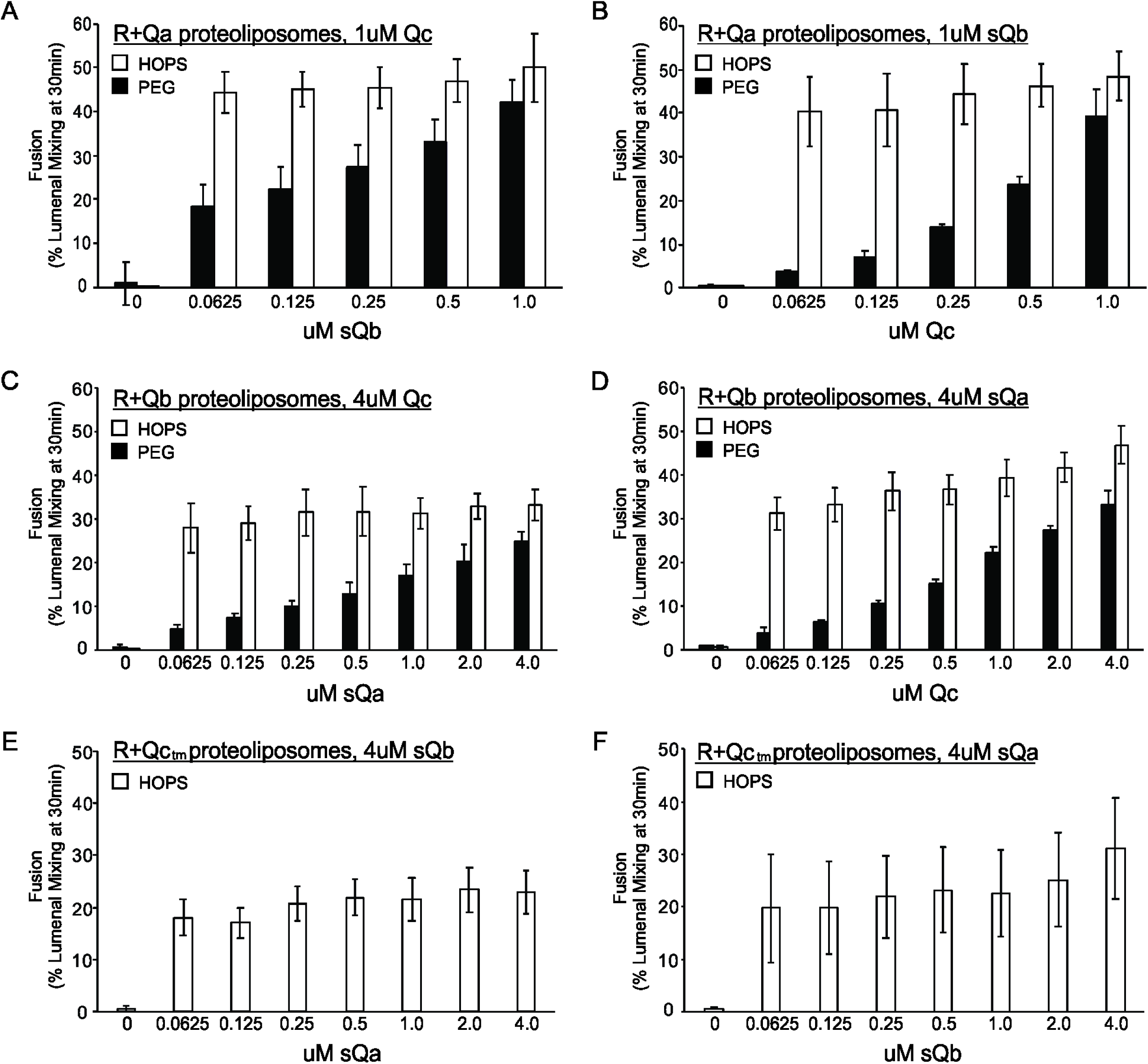
Fusion with HOPS is saturable for each vacuolar Q-SNARE. Reconstituted proteoliposomes of VML composition were prepared with wild-type Ypt7 at 1:4000 protein to lipid molar ratio and either R-SNARE or each single Q-SNARE at a 1:2500 protein to lipid ratio, employing a transmembrane version of Qc. Fusion assays were performed in RB150. Ypt7/R-SNARE proteoliposomes and Ypt7/Q-SNARE proteoliposomes were separately incubated at 1mM (lipid) with 20µM streptavidin, 2mM EDTA, 0.5mM MgCl_2_, and 1mM GTP for 10 min at 27°C. MgCl_2_ was then added to bring the concentration to 2.5mM. The nucleotide-exchanged R- and Q-proteoliposomes were then combined and portions were added to tubes containing one half volume of either 0.16µM HOPS or 8% PEG. Aliquots of each (16µl) were pipetted into a 384 well plate. During the nucleotide exchange process, a mixture of the missing soluble Q-SNAREs was prepared in RB150, containing 4µM of each soluble Q-SNARE (A and B) or 16µM of each soluble Q-SNARE (C, D, E, and F). Two dilution curves were then prepared, keeping one soluble SNARE at the starting concentration while diluting the other 2-fold. A portion (5µl) of each dilution was pipetted into empty wells of a 384 well plate, which then received 15µl of the mixtures of proteoliposome with HOPS or PEG. Final concentrations of HOPS or PEG in the 20µl reaction were 40nM and 2%, respectively.

## Discussion

The proteins and lipids which mediate homotypic vacuole fusion cluster around the edge of the apposed membranes of docked vacuoles (Wang et al., 2002) and are interdependent for this localization (Fratti et al., 2004). The multiplicity of affinities among these proteins is striking, and may underlie the interdependent character of their microdomain enrichment and functions for fusion. For example, HOPS binds each of the 4 SNAREs (Stroupe et al., 2006; Baker et al., 2015; Figure 4), acidic lipids (Karunakaran and Wickner, 2013), phosphoinositides (Stroupe et al., 2006), Ypt7 on each docked membrane (Hickey and Wickner, 2010), and Sec17 (unpublished). While multiple binding affinities are perhaps expected for a large, multi-subunit complex such as HOPS, even Sec17 binds to SNAREs (Söllner et al., 1993; Zick et al., 2015), lipid (Zick et al., 2015), Sec18 (Weidman et al., 1989), HOPS (Figure 8), and itself (T. Torng and W. Wickner, unpublished). Each of these components is required for fusion, both *in vivo* and *in vitro* with purified vacuoles. The purification of each of these proteins allows exploration of their mutual affinities, while the creation of natural and synthetic fusion sub-reactions allows tests of their functionality.

Though HOPS recognizes each SNARE, the functions of these recognitions are not fully resolved. HOPS may catalyze SNARE assembly or integrate SNARE functions with lipid and SNARE chaperones. In one model sub-reaction (Orr et al., 2017), proteoliposomes bearing Ypt7 and R-SNARE were incubated with the 3 soluble Q-SNAREs (lacking transmembrane anchor) and HOPS, then re-isolated by floatation. HOPS was required for the association of each of the Q-SNAREs, and each Q-SNARE depended on the other two for its HOPS-dependent membrane association. This is a clear case of HOPS-dependent 4-SNARE complex assembly. In other studies, a synthetic tether can substitute for HOPS to allow efficient functional *trans*-SNARE complexes to assemble between 3Q- and R-proteoliposomes and lead to fusion (Song and Wickner, 2019). When the Q-SNARE fusion partner has only two anchored Q-SNAREs instead of all three, HOPS is essential for fusion, and GST-PX will not suffice (Figure 1). HOPS can catalyze the entry of each Q-SNARE into fusion-competent complex (Figures 1, 3).

The current findings suggest an update to the model of SM protein function. With the discovery (Baker et al., 2015) of conserved grooves on the surface of the HOPS SM-family subunit Vps33 which bind the R- and Qa-SNARE domains in parallel (N to C) and in register (with adjacent 0-layer residues), it was proposed that these associations are the key committed step for 4-SNARE assembly. However, it is now clear that assembly of R-with Q-SNAREs proceeds without SM function as long as there is tethering and the three Q-SNAREs are pre-assembled. HOPS has the unique capacity to create a fusion-ready assembly of proteoliposomes bearing R-SNARE with those bearing any two Q-SNAREs, able to receive the third Q-SNARE for rapid fusion.

It remains unclear whether all the components needed for fusion remain engaged with each other up to and during lipid bilayer mixing; is there a 2Ypt7/HOPS/4SNARE/2Sec17/Sec18 complex? Earlier studies of *cis*-SNARE complexes from isolated vacuoles showed that HOPS and Sec17 were in separate complexes with SNAREs, and suggested that Sec17 could displace HOPS from SNARE associations (Collins et al., 2005). It is also unclear whether HOPS remains bound to Ypt7 and even whether it remains bound to the SNAREs. We have noted (Baker et al., 2015) that the helical R and Qa SNARE domains bind to their sites on Vps33, the HOPS SM-family subunit, with the same face of the SNARE domain helix in contact with Vps33 as faces inward toward the other SNAREs in the assembled 4-helical SNARE complex. These SNAREs are thus likely to leave their Vps33 contact sometime prior to completion of SNARE zippering, though the whole SNARE complex may exploit the HOPS affinities for the Qb and Qc SNAREs to remain bound.

The vacuole fusion reaction has been studied in cells, with the isolated organelle, and with purified components reconstituted into proteoliposomes. The latter approach allows reconstitution and assay of subreactions, addressing mechanistic questions and testing and revising models. In early models of fusion, tethering provides SNAREs proximity for *trans*-complex assembly, either spontaneous or catalytically initiated by R- and Qa-SNARE domain templating on an SM protein. SNAREs then zippered spontaneously, distorting the bilayers for fusion. After fusion, SNARE NSF/Sec18 and αSNAP/Sec17 function as an ATP-driven SNARE disassembly chaperone system to disassemble *cis*-SNARE complexes for the subsequent round of fusion. Recent mechanistic studies have refined this model. Tethering brings all the fusion proteins and lipids into proximity, allowing an interdependent enrichment in a dedicated fusion microdomain. Membrane tethering also allows SNAREs to assemble in *trans* in a fusion-competent conformation (Song and Wickner, 2019). Large Rab-effector complexes, such as vacuolar/lysosomal HOPS, will mediate tethering (Baker and Hughson, 2016) and, along with SM proteins, may guide SNARE complex assembly. HOPS coordinates the loading of Sec17 and Sec18 onto SNAREs, promoting fusion by some combination of adding wedge-like bulk to the fusion domain (Agostino et al., 2017), promoting SNARE zippering (Song et al., 2017), and distorting bilayers adjacent to the SNAREs with the Sec17 apolar loop (ibid). It remains unclear whether HOPS and Sec17 remain associated with *trans*-SNARE complexes at the same time and how Sec18 can contribute to fusion without disassembling the *trans*-SNARE complexes.

## Materials and Methods

### Proteins and reagents

The soluble version of GST-Qa (GST-sQa), with Qa amino acyl residues 1-264 but lacking its transmembrane domain, was generated by PCR with the Phusion high-fidelity DNA polymerase (NEB). The DNA fragment was cloned into BamHI and SalI digested pGST parallel1 vector (Sheffield et al., 1999) with an in-Fusion kit (Clonetech).

For GST-sVam3,

F: AGGGCGCCATGGATCCGATGTCCTTTTTCGACATCGA

R: AGTTGAGCTCGTCGACTACTTACCGCATTTGTTACGGT

Trans-membrane (tm)-anchored Ypt7: The nucleotide sequence encoding the transmembrane domain of the Qa-SNARE Vam3 (amino acyl residues 265-283) fused to the 3’ end of the nucleotide sequence encoding full length Ypt7 was amplified by PCR from pET-19 Ypt7-tm (a kind gift from C Ungermann) with the Phusion high-fidelity DNA polymerase (NEB). The DNA fragment was cloned into BamHI and SalI digested pMBP-parallel1 vector (Sheffield et al., 1999), with the HiFi DNA assembly kit (New England Biolabs, Ipswich, MA).

For Ypt7-tm

F: AGGGCGCCATGGATCCGTCTTCTAGAAAAAAAAATATTTT

R: AGTTGAGCTCGTCGACTAACTTAATACAGCAAGCA

The resulting plasmid sequence was confirmed.

The purifications of HOPS (Zick and Wickner, 2013), GST-PX (Fratti et al., 2004), Sec17p (Schwartz and Merz, 2009), Sec18p (Mayer et al., 1996), wild-type Ypt7 (Zick and Wickner, 2013), and a soluble version of MBP-Qb (MBP-sQb) lacking its transmembrane domain (Zick and Wickner, 2013) were as described. Full-length, wild-type vacuolar SNAREs GST-Qa, Qc, R, and Qb were isolated as described (Mima et al., 2008; Schwartz and Merz, 2009; Zucchi and Zick, 2011), and Qb and R were buffer exchanged into β-octylglucoside (Zucchi and Zick, 2011). Vam7-tm (Xu and Wickner, 2012) and Sec17-tm (Song et al., 2017) were purified as described. The plasmid encoding His_6_-Vam3 (full length) was a kind gift from Joji Mima, and the protein was purified as described (Izawa et al., 2012).

MBP-Ypt7-tm was purified as follows: MBP-Ypt7-tm was produced in *E.coli* Rosetta(DE3)*pLysS* (Novagen, Milwaukee WI). A single colony was inoculated into 50ml LB medium containing 100 μg/ml ampicillin (Amp) and 37 μg/ml Chloramphenicol (Cam) and grown overnight at 37°C, then transferred to 6 liters LB with 100 μg/ml Amp and 37 μg/ml Cam. Cultures were grown at 37°C to an OD_600_ of 0.5. IPTG (0.5 mM) was added and cultures were shaken for 3 hr at 37°C. Cells were harvested by centrifugation (Beckman JA10 rotor, 5000rpm, 5min, 4°C) and resuspended in 50 ml buffer A (20 mM HEPES/NaOH, pH 7.4, 100 mM NaCl, 1 mM EDTA, 1 mM DTT, 1 mM PMSF (phenylmethylsulfonyl fluoride) and PIC (protease inhibitor cocktail; (Xu and Wickner, 1996)). Resuspended cells were lysed by French Press (8,000 psi, 4°C, 2 passages) and lysates were centrifuged (Beckman 60Ti rotor, 30 min, 50,000 rpm, 4°C). Pellets were resuspended in 100 ml of buffer B (PBS [140 mM NaCl, 2.7 mM KCl, 10 mM Na_2_HPO_4_ and 1.8 mM KH_2_PO_4_, pH7.4], 1 mM EDTA, 1 mM dithiothreitol, 10% glycerol, PIC and 1 mM PMSF) with a Dounce homogenizer and centrifuged (60Ti, 50,000 rpm, 30 min, 4°C). Pellets were resuspended in 100 ml of buffer C (PBS, 1 mM EDTA, 1 mM DTT, 1% Triton X100, 10% glycerol, PIC and 1 mM PMSF) with a Dounce homogenizer and incubated (4°C, 30 min) with nutation for 1hr. The extract was centrifuged (60Ti, 50,000 rpm, 30 min, 4°C) and the supernatant was added to 24 mL of amylose resin (NEB, Ipswich MA) pre-equilibrated with buffer C and nutated for 2 hr at 4°C. The resin was gravity-packed into a 2.5 cm diameter column at 4°C, washed with 100 mL of buffer D (100 mM HEPES/NaOH, pH 7.4, 100 mM NaCl, 1 mM EDTA, 1 mM DTT, 100 mM *β*-OG, 10% glycerol). MBP-Ypt7-tm was eluted with 25 mM maltose in buffer D. Proteins were frozen in liquid nitrogen and stored at −80°C.

A plasmid encoding the soluble version of GST-R (GST-sR) lacking its transmembrane domain (Thorngren et al., 2004) was transformed into *E.coli* BL21(DE3) and the protein was purified as follows: 100ml of LB+ 100µg/ml Ampicillin was inoculated with a single colony, shaken overnight at 37°C, then added to 3L of LB+Ampicillin. Cultures were grown at 37°C to an OD_600_ of 0.8, induced with 1mM IPTG, and shaken overnight at 18°C. Cells were harvested and resuspended in 40mls resuspension buffer (20mM TrisCl, pH 8.0, 200mM NaCl, 200µM PMSF, PIC). Cells were lysed by French Press (2 passages) and lysates were centrifuged in a Beckman 60ti rotor (1hr, 50,000 rpm, 4°C). The supernatant was nutated (2h, 4°C) with 10ml glutathione-Sepharose 4B resin (GE Healthcare, Pittsburg, PA) in resuspension buffer. The slurry was poured into a column, the settled resin was washed with resuspension buffer, and protein eluted with 100mM HEPES-NaOH pH 7.8, 300mM NaCl, 20mM glutathione. The protein peak was dialyzed into RB150 (20mM HEPES-NaOH pH 7.4, 150mM NaCl, 10% glycerol [vol/vol]) in 6-8K molecular weight cutoff dialysis tubing (Fisher Scientific, Pittsburgh, PA), aliquoted, and frozen in liquid nitrogen. GST-sVam3 was purified the same way as GST-sNyv1, except that the growth media also contained 37 μg/ml chloramphenicol, the culture was grown to OD_600_ of 1.0 before induction, the elution buffer was 20mM HEPES-NaOH pH 7.4, 300mM NaCl, 20mM glutathione, 1mM DTT, and the eluate was frozen in aliquots without dialysis. Before use, the MBP-sVti1, GST-sNyv1, and GST-sVam3 were cleaved with TEV protease to remove their tags, unless otherwise noted.

Dilinoleoyl lipids (diC18:2 PC, PS, PE, and PA), soy PI, and 1,2-dipalmitoyl-*sn*-glycerol were purchased from Avanti Polar Lipids (Alabaster, AL). Ergosterol was from Sigma Aldrich (St. Louis, MO), PI(3)P from Echelon Biosciences (Salt Lake City, UT), and the fluorescent lipids Marina-Blue DHPE, NBD-PE, and Lissamine rhodamine DHPE were from Invitrogen by Life Technologies (Eugene, OR). N-octyl-ß-D-glucopyranoside was from Anatrace (Maumee, OH), and poly(ethylene glycol) 8,000 was from Sigma-Aldrich.

### Proteoliposome preparation

Proteoliposomes were prepared as described in Zick et al. (2014) with modifications. Lipid compositions of vacuolar mimic lipid (VML) proteoliposomes for content-mixing assays were 47.3 or 46.1 mol% diC18:2 PC, 18% diC18:2 PE, 18% soy PI, 4.4% diC18:2 PS, 2% diC18:2 PA, 8% ergosterol, 1% diacylglycerol, 1% diC16 PI(3)P and either 0.3% Marina Blue-PE or 1.5% NBD-PE. Lipid compositions of proteoliposomes for flotation assays were either 99% diC18:2 PC and 1% Lissamine rhodamine-DHPE or % diC18:2 PC, 15% diC18:2 PS and 1.5% NBD-PE. Proteins were added at protein:lipid ratios as described in the figure legends. Proteoliposomes were isolated by flotation through density medium as described (Zick and Wickner, 2013) and assayed for total phosphate (Chen et al., 1956). Aliquots of proteoliposomes in RB150+Mg^2+^ (20mM HEPES-NaOH, pH 7.4, 150mM NaCl, 10% glycerol [vol/vol], 1mM MgCl_2_) were frozen in liquid nitrogen at a concentration of 2mM lipid phosphorus.

### Flotation assay

Flotation assays were conducted as described (Orr et al., 2017) with modifications. Liposomes (7.5µl) were incubated for 1 h at 30°C in 30µl reactions (0.5mM lipid, 500nM HOPS, 0.2% defatted bovine serum albumin (BSA; Sigma-Aldrich), and 1mM MgCl_2_ in RB150). Reactions were gently vortexed with 90µl of 54% (wt/vol) Histodenz (Sigma-Aldrich) in iso-osmolar RB150/Mg^2+^ (containing a reduced level (2%) of glycerol) and 80µl were transferred to 7 × 20 mm polycarbonate tubes (Beckman Coulter, Brea CA), overlaid with 80µl of 35%, then 80µl of 30% Histodenz in iso-osmolar RB150+Mg^2+^ and finally 50µl of RB150+Mg^2+^. The remaining portions of the starting incubations were solubilized with 1µl of 5% (vol/vol) Thesit for determination of lipid recovery. Reactions were centrifuged in a Beckman TLS-55 rotor, 4°C, 55,000 rpm, 30 min. Samples were harvested by pipetting 80µl from the top of the tube and solubilized with 2µl of 5% Thesit. Lipid recovery was assayed as described (Orr et al., 2014), measuring either rhodamine fluorescence (excitation, 560nm; emission, 580nm; cutoff 570) or NBD fluorescence (excitation, 460nm; emission, 538nm; cutoff 515), depending on the composition of the liposomes. Bound protein determination was performed as described (Orr et al., 2014).

### Fusion assay

Proteoliposomes were nucleotide exchanged by incubating proteoliposomes (1 mM lipid), RB150, streptavidin (10 μM), EDTA (2 mM), and GTP (20 μM) for 10 min at 27°C. Nucleotide exchange was completed by adding MgCl_2_ (4 mM) and the mixture was placed on ice. The fusion reaction was initiated by mixing 5 μl each of GTP exchanged R- and Q-SNARE proteoliposomes with soluble components (10 μL of e.g., HOPS, GST-PX, and soluble SNAREs as noted), for a total volume of 20ul. Plates (Corning 4514, 384 wells) were incubated at 27°C in SpectraMax Gemini XPS (Molecular Devices, Sunnyvale, CA) fluorescence plate reader and lumenal mixing was assayed every minute, as described (Zick and Wickner, 2016).

## Acknowledgements

We thank Michael Zick, Jose Rizo, Gustav Lienhard, and Charles Barlowe for fruitful discussions, and Christian Ungermann for the generous gift of a plasmid encoding Ypt7-tm. This work was supported by NIH grant R35GM118037. MH was supported by Deutsche Forschungsgemeinschaft fellowship HA 7730/2-1.

## Figure Supplements

**Figure 1-figure supplement 1.**
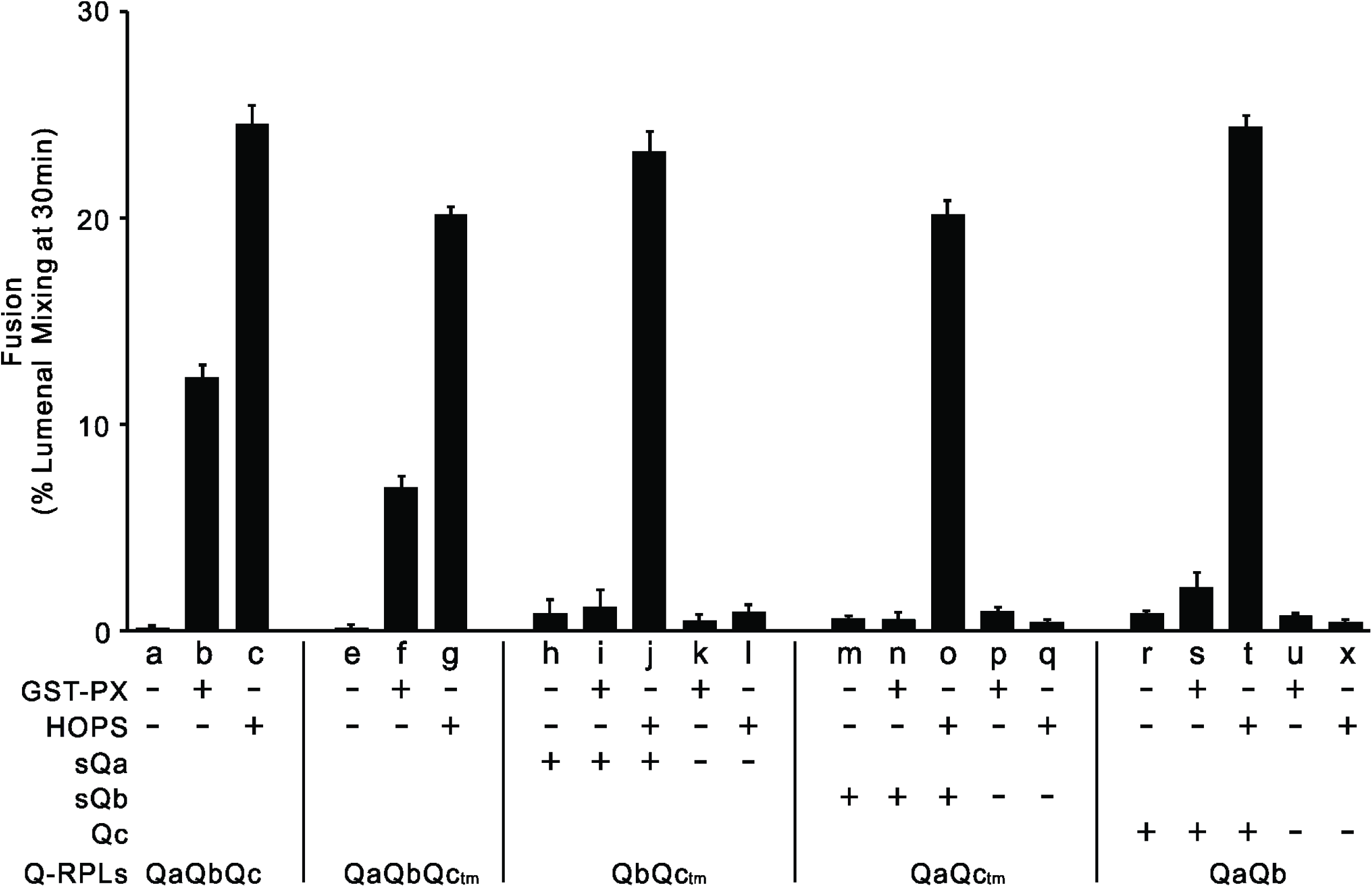
HOPS, but not GST-PX, supports fusion of R- and 2Q-SNARE proteoliposomes mixed with the third soluble Q-SNARE. Fusion assays were conducted as described in Figure 1, with R and QaQbQc, QaQbQc_tm_, QbQc_tm_, QaQc_tm_ or QaQb proteoliposomes. All proteoliposomes had SNAREs at a 1:16000 and Ypt7-tm at a 1:8,000 protein: lipid molar ratio. Average and standard deviations of fusion 30 min after addition of HOPS, for triplicate assays relative to total mixed contents.

**Figure1-figure supplement 2.**
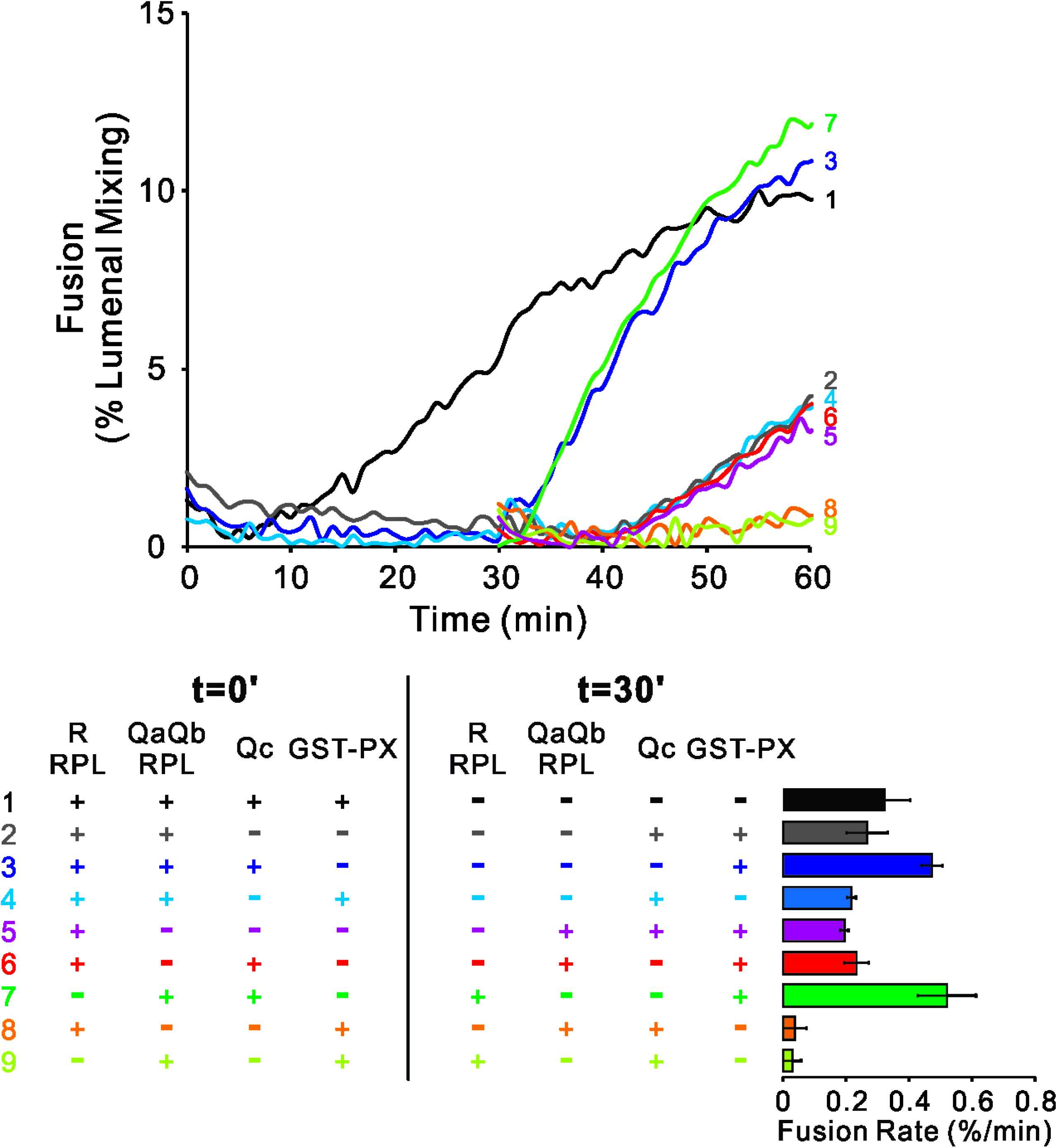
The Qc-SNARE can slowly and stably assemble spontaneously with QaQb-proteoliposomes. Fusion reactions had proteoliposomes bearing either the R- or QaQb-SNAREs at 1:16000 SNARE:lipid molar ratio. R and QaQb proteoliposomes were mixed without preincubation (1,2,3 and 4) or after 30 min preincubation at 27°C (5,6,7,8 and 9). Fusion incubations received 500nM GST-PX and/or 4μM Qc where indicated. The bar graph quantifies the maximal rate of content mixing from three independent experiments. The rate of fusion was determined as the slope for 10min of the content mixing reaction after fusion initiation. Kinetic curves of contents mixing assays in this figure are representative of n ≥ 3 experiments.

**Figure 1-figure supplement 3.**
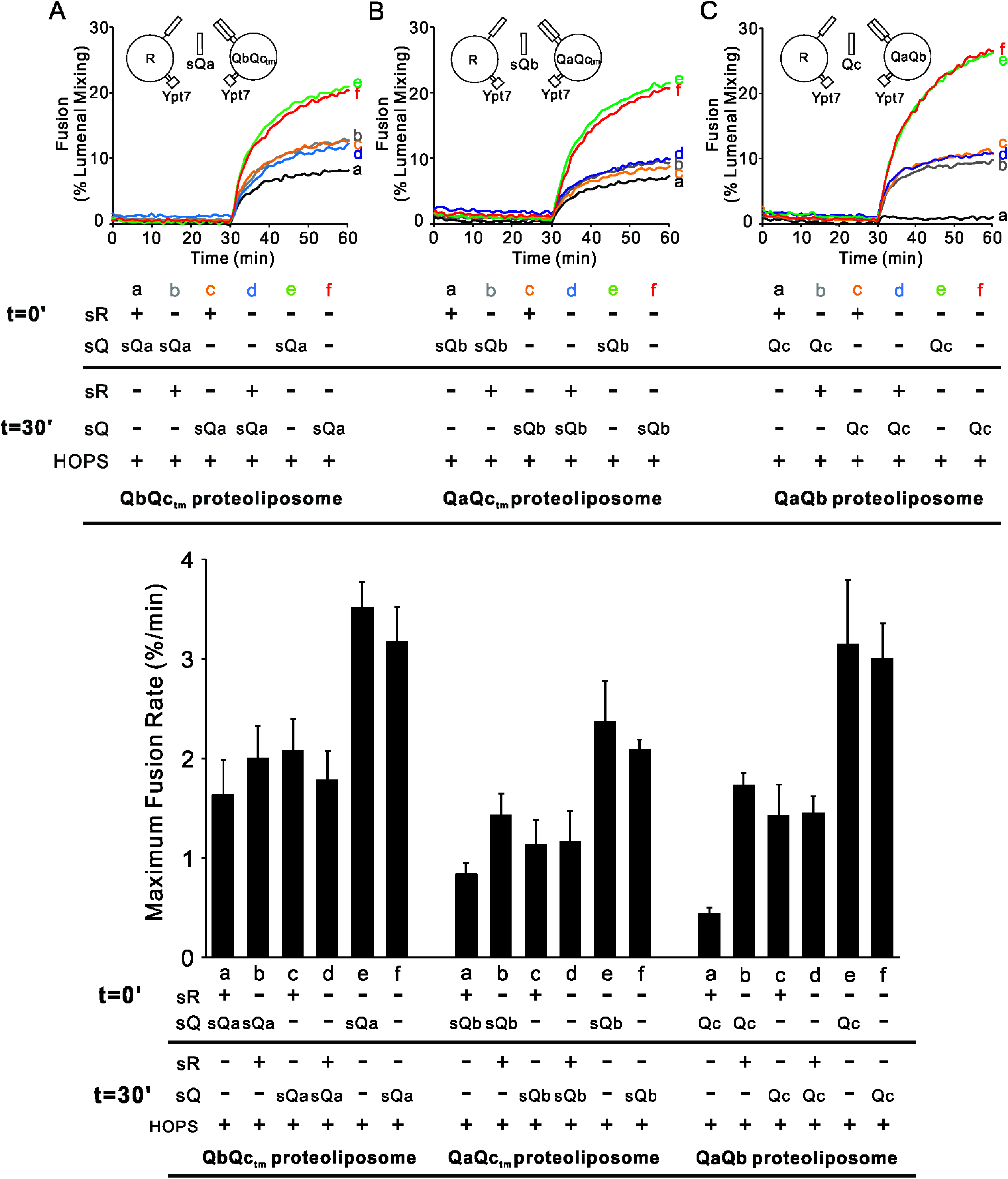
Fusion inhibition by sR. Fusion reactions had proteoliposomes bearing either the R or (A) QbQc_tm_, (B) QaQc_tm_, or (C) QaQb SNAREs and Ypt7-tm at 1:16000 SNARE:lipid and 1:8000 Ypt7:lipid molar ratios. Fusion incubations received 50 nM HOPS at t=30 with 4μM of the soluble form of the third Q-SNARE at t=0 (a, b, e) or t=30 (c, d, f) as indicated. Soluble Nyv1 (sR) was added to 4μM at t=0 (a, c) or t=30 (b, d). Content mixing assays in this figure are representative of n ≥ 3 experiments; means and standard deviations for each experiment are presented.

**Figure 3-figure supplement 1.**
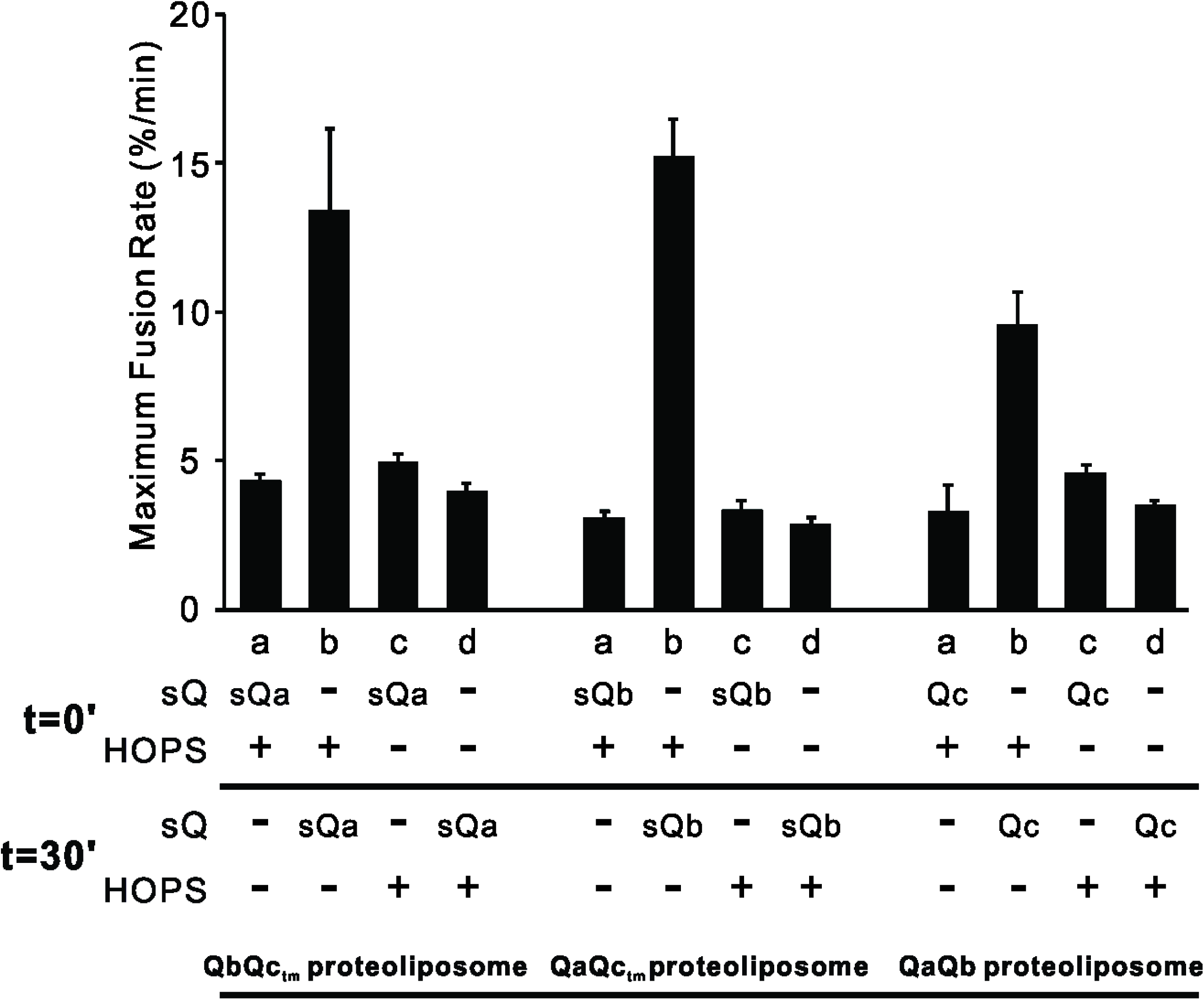
Preincubation of R- and 2Q-SNARE proteoliposomes with HOPS gives more rapid fusion when the third soluble SNARE is added than when all components are mixed without preincubation. Fusion assays were as described in Figure 3, with R and QbQc_tm_, QaQc_tm_ or QaQb SNARE proteoliposomes. All proteoliposomes had SNAREs at a 1:16000 protein: lipid molar ratio and Ypt7-tm at a 1:8,000 protein: lipid molar ratio. The maximum rates of fusion were determined. Average and standard deviations of fusion rate from three independent experiments.

**Figure 4-figure supplement 1.**
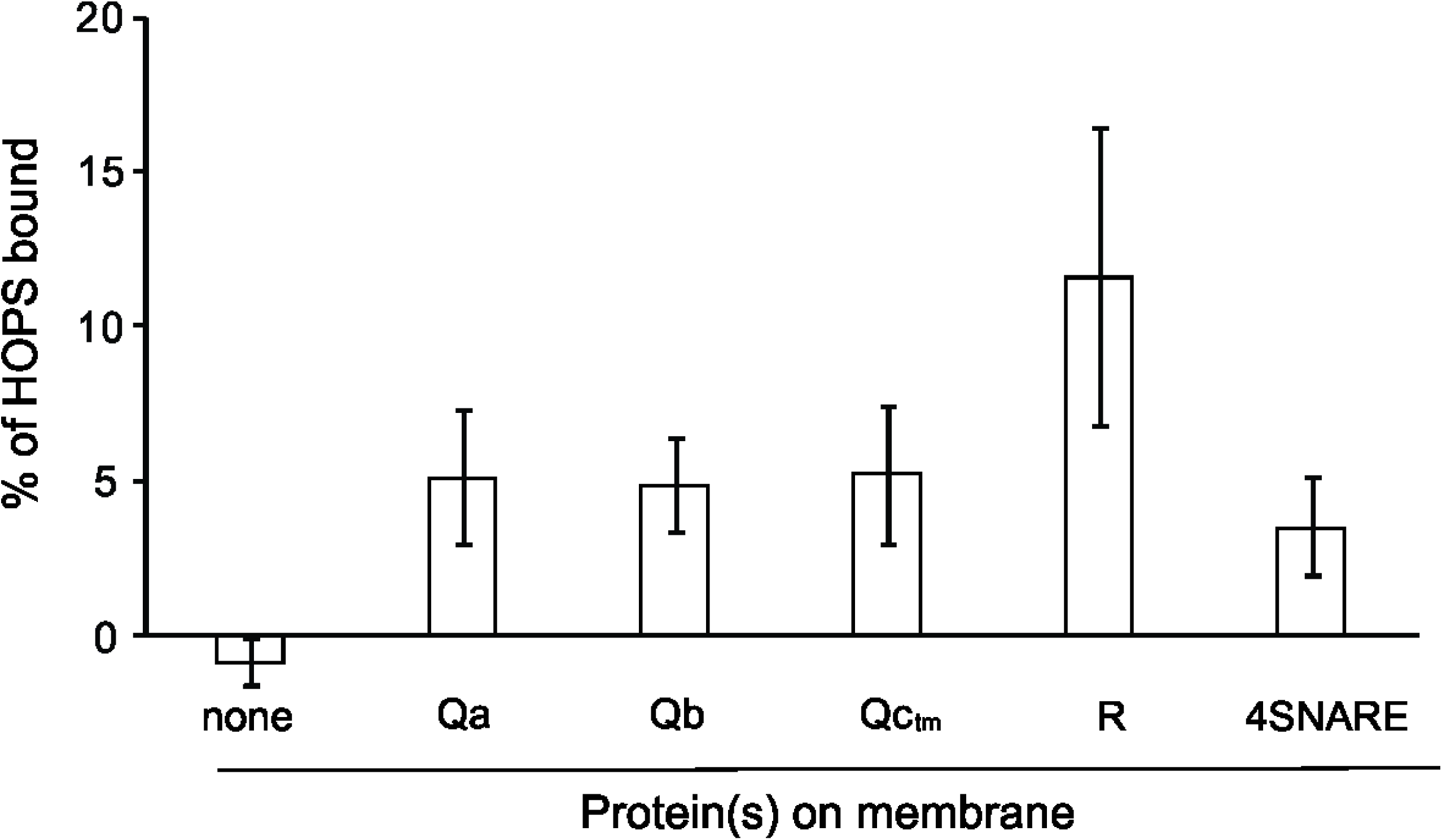
HOPS reproducibly binds to each SNARE. Western blots of 3 independent experiments were analyzed with UN-SCAN-IT software (Silk Scientific, Orem UT).

**Figure 5-figure supplement 1.**
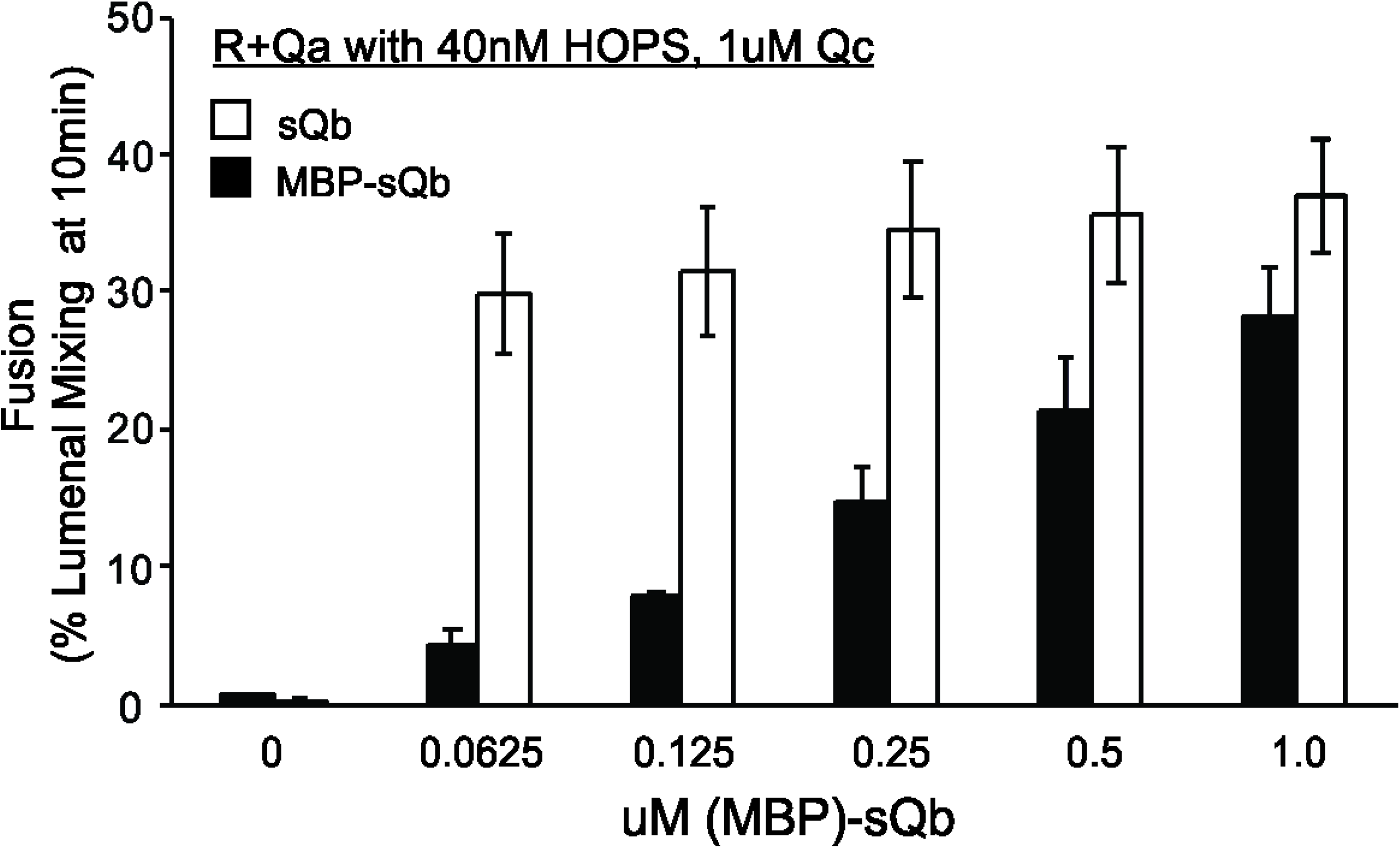
An MBP tag on the soluble Qb-SNARE interferes with its recruitment by HOPS. Fusion assays were conducted as described in Figure 5, using R-SNARE+Ypt7 and Qa-SNARE+Ypt7 RPLs, 40nM HOPS, a constant (1µM) level of Qc, and 2-fold decreasing concentrations of MBP-tagged soluble Qb that either had or had not been cleaved to remove its MBP domain by a 2 hour incubation with TEV protease.

## Source Data

**Figure 1-Source Data 1.** Source data file (Excel) for Figure 1A, B, C, D and E.

**Figure 2-Source Data 1.** Source data file (Excel) for Figure 2A and C.

**Figure 3-Source Data 1.** Source data file (Excel) for Figure 3A, B and C.

**Figure 5-Source Data 1.** Source data file (Excel) for Figure 5A and B.

**Figure 5-Source Data 2.** Source data file (Excel) for Figure 5C and D.

**Figure 5-Source Data 3.** Source data file (Excel) for Figure 5E and F.

